## Supplementary Information for "Ultrathin Liquid Cells for Microsecond Time-Resolved Cryo-EM"

**This PDF file includes:**

1 | Sample preparation

2 | Deposition of sealing layers onto cryo samples

3 | Laser melting and revitrification experiments

4 | Data collection and analysis – apoferritin

5 | Data collection and analysis – 50S ribosomal subunit

6 | Cryo-EM data collection and processing statistics

7 | Irradiation with long laser pulses and breakup of the thin liquid film

8 | Variability analysis of the L1 stalk motion of the 50S ribosomal subunit

9 | Determination of the conformational distributions for the L1 stalk movement

10 | Simulation of the sample temperature during laser heating

11 | Estimate of the particle temperature during laser heating

12 | Evolution of the amplitude of the L1 stalk motion

**1 | Sample preparation**

Mouse heavy chain apoferritin (8 mg/ml, 10 mM HEPES buffer, pH 7.5, 150 mM sodium chloride) was provided by the Protein Production and Structure Core Facility at EPFL, and the 50S ribosomal subunit (40 $OD_{260}$/ml, 20 mM HEPES buffer, pH 7.5, 100 mM sodium chloride, 2 mM magnesium chloride) by Dr. Bertrand Beckert of the Dubochet Center for Imaging in Lausanne.[1] Cryo samples are prepared by applying 3 µl of the sample solution onto UltrAuFoil grids (R1.2/1.3, 300 gold mesh, Quantifoil) that were plasma cleaned for 90 s to render them hydrophilic (EasyGlow, TedPella, air). The samples are plunge-frozen with a Thermo Fisher Vitrobot Mark IV (4 °C, 95% relative humidity, blotting force 10, 6 s blotting time for apoferritin; 20 °C, 95% relative humidity, blotting force 10, 2 s blotting time for the 50S ribosomal subunit).

**2 | Deposition of sealing layers onto cryo samples**

A custom-built vapor deposition setup is used to deposit ultrathin silicon dioxide layers onto both sides of the cryo sample. The samples are placed in a single tilt cryo specimen holder (Elsa, Gatan), which is inserted into the deposition chamber through a load-lock that was repurposed from a retired JEOL transmission electron microscope. Silicon dioxide is vapor deposited using an effusion cell (TecTra e-flux) at a constant deposition rate of 0.5 Å/s. The cryo holder is first rotated such that the top surface of the sample faces the effusive beam, and the top sealing layer is deposited, after which the holder is rotated again to deposit the bottom sealing layer. The thickness of the deposited membranes (1.4 ± 0.1 nm) is determined with a quartz crystal microbalance[2] onto which the effusive beam is simultaneously deposited.

**3 | Laser melting and revitrification experiments**

Revitrification experiments are performed as described previously, using a modified JEOL 2200FS transmission electron microscope.[3,4] Microsecond laser pulses are generated by chopping the output of a continuous wave laser (Novanta Laser Quantum Ventus, 532 nm wavelength) with an acousto-optic modulator (AA Opto-Electronic). We typically use a laser power of about 80 mW and estimate that we have losses of at least a quarter between the point at which we measure the laser power and the sample. The laser beam is focused to a spot size of 28 ± 2 µm FWHM in the sample plane for apoferritin and 22 ± 2 µm for the 50S ribosomal subunit, as determined with a camera that is placed in a plane conjugate to the sample plane. Melting and revitrification experiments are performed by aiming the laser beam onto the center of a grid square. The laser power is adjusted such that a single 30 μs laser pulse revitrifies an area of about 9–25 holes. Apoferritin cryo samples are revitrified with a train of 6 laser pulses of 35 μs duration, and samples of the 50S ribosomal subunit with 1, 5, or 10 laser pulses of 30 μs duration (about 10–15 ms apart).

**4 | Data collection and analysis – apoferritin**

High-resolution micrographs of a conventional cryo sample of apoferritin as well as a sample that was sealed and revitrified were collected at the Dubochet Center for Imaging in Lausanne, using a Thermo Fisher Titan Krios G4 equipped with a Falcon 4i camera and a SelectrisX energy filter. The data acquisition parameters are summarized in Supplementary Table 1. Single-particle reconstructions were performed in CryoSPARC v4.4.1,[5] with the corresponding workflows illustrated in Figs. S1 and S2.

The micrographs were patch motion corrected, and movies with a total full-frame motion distance of more than 50 pixels were discarded. Contrast transfer function (CTF) estimation was performed using Patch CTF in CryoSPARC. Micrographs with an estimated resolution of worse than 6 Å or an astigmatism of over 1,000 Å were discarded, as well as micrographs that showed hexagonal or cubic ice upon inspection. Approximately 1,000 particles were first picked manually in each data sets and subjected to 2D classification. The 2D classes were then used for template picking (120 Å diameter). The particles were extracted with a box size of 440 pixels. Following 2D classification, *ab initio* reconstruction ($C_1$ symmetry), and homogeneous refinement (O symmetry), all the extracted particles were subjected to three rounds of heterogeneous refinement (O symmetry), using 5 decoy classes and the homogeneously refined map as initial volumes. The best class resulting from the heterogeneous refinement was further refined with octahedral symmetry imposed, using non-uniform refinement, global and local CTF refinement, per particle defocus refinement, and Ewald sphere correction. From the conventional sample, 170,473 particles were randomly selected to match the number of particles obtained for the revitrified sample. The global and local resolution was estimated using a gold-standard FSC threshold of 0.143. The structures were visualized with UCSF ChimeraX 1.7.[6,7]

Apoferritin – Conventional

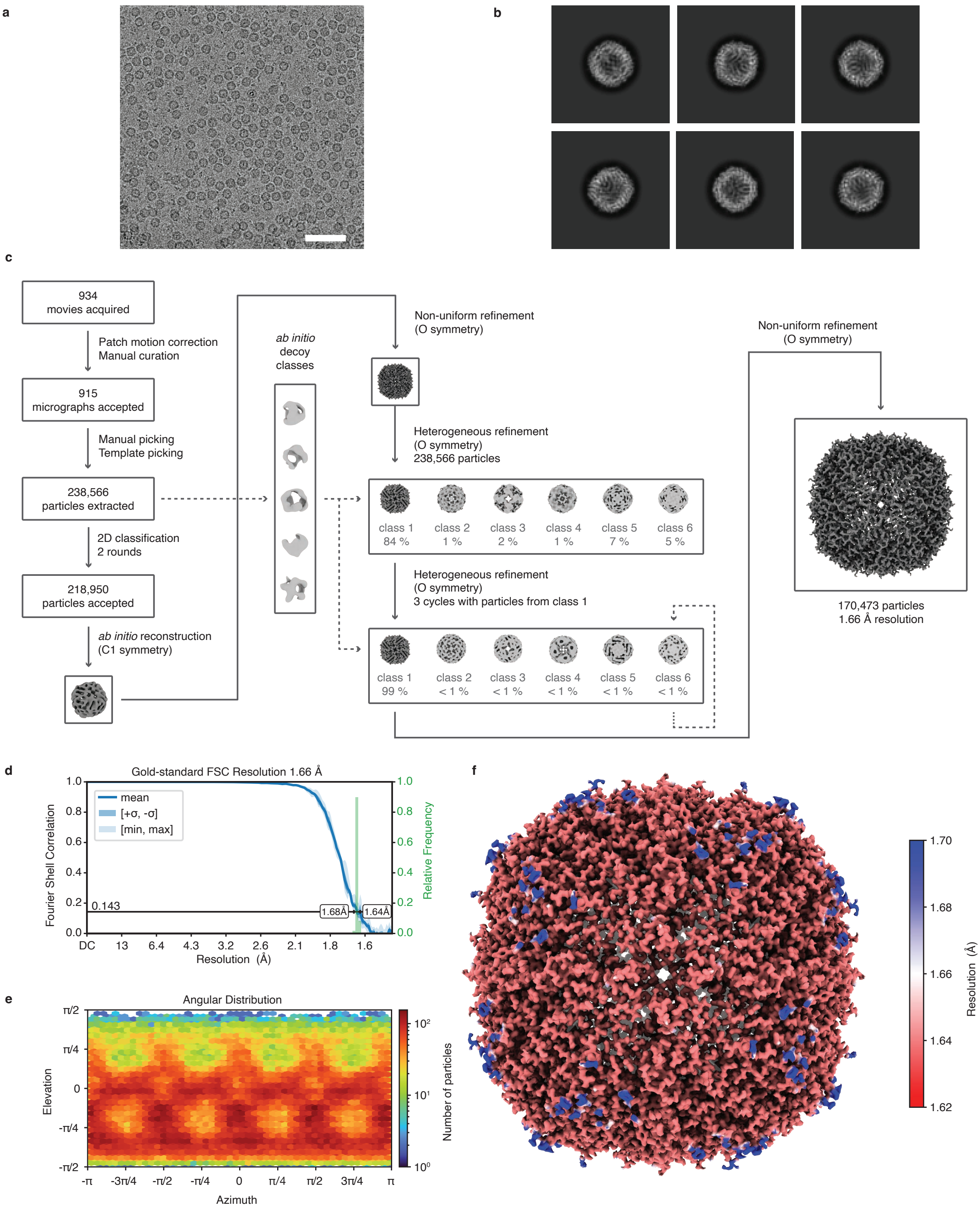

**Figure S1 | Cryo-EM processing workflow for the conventional cryo sample of apoferritin.** **a,** Representative micrograph. Scale bar, 50 nm. **b,** 2D class averages gathered from the processing workflow. **c,** Data processing workflow in CryoSPARC. The symmetry applied is indicated in parentheses. The map is shown at a threshold of 5 σ above the mean. **d,** Conical FSC, with the cutoff of 0.143 indicated by a black line. The mean FSC is shown as a blue line, with the dark blue shading indicating one standard deviation, and the light blue shading the minimum and maximum values of the conical FSC. A histogram of the resolution values is shown in green, with the minimum and maximum values indicated. **e,** Orientation distribution of the particles. **f,** Final map with the local resolution estimation indicated in color (GSFSC cutoff of 0.5).

**Apoferritin – sealed and revitrified, 210 μs**

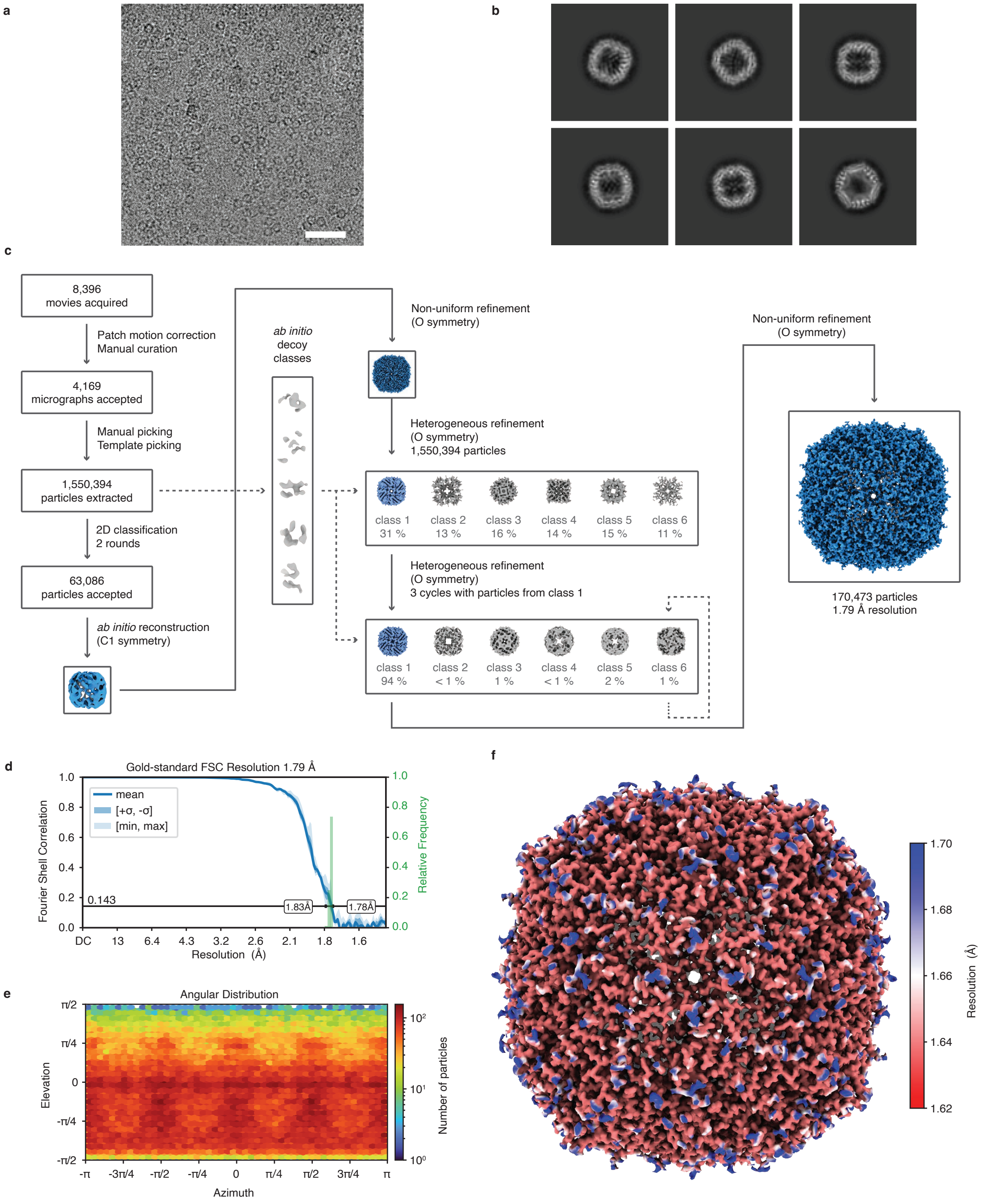

**Figure S2 | Cryo-EM processing workflow for the sealed and revitrified apoferritin specimen.** **a,** Representative micrograph. Scale bar, 50 nm. **b,** 2D class averages gathered from the processing workflow. **c,** Data processing workflow in CryoSPARC. The symmetry applied is indicated in parentheses. The map is shown at a threshold of 5 σ above the mean. **d,** Conical FSC, with the cutoff of 0.143 indicated by a black line. The mean FSC is shown as a blue line, with the dark blue shading indicating one standard deviation, and the light blue shading the minimum and maximum values of the conical FSC. A histogram of the resolution values is shown in green, with the minimum and maximum values indicated. **e,** Orientation distribution of the particles. **f,** Final map with the local resolution estimation indicated in color (GSFSC cutoff of 0.5).

**5 | Data collection and analysis – 50S ribosomal subunit**

High-resolution data of all 50S Ribosome samples were collected at Dubochet Center for Imaging, Lausanne, Switzerland (DCI-Lausanne) using a Thermo Fisher Titan Krios G4 microscope equipped with Falcon 4i camera and SelectrisX energy filter. The data acquisition parameters are given in Supplementary Table 2. Single-particle reconstructions were performed in CryoSPARC 4.4.1,[5] with the corresponding workflows illustrated in Figs. S3–S6.

The micrographs were patch motion corrected, and movies with a total full-frame motion distance of more than 50 pixels were discarded. Contrast transfer function estimation was performed using Patch CTF in CryoSPARC. Micrographs with an estimated resolution of worse than 8 Å or an astigmatism of over 1,000 Å were discarded, as well as micrographs containing hexagonal or cubic ice upon inspection. Particles from the conventional, the 30 μs, and the 150 μs data sets were picked using a blob picker with a diameter ranging from 200 to 300 Å. The particles were extracted with a box size of 784 px and Fourier-cropped to 392 px. For the 300 μs data set, all accepted micrographs were denoised using the Denoiser job in CryoSPARC, which was trained on 100 randomly selected micrographs. Particles were first picked using a blob picker with a diameter of 200–300 Å, then sorted using two rounds of 2D classification, followed by *ab initio* reconstruction and homogeneous refinement. The obtained volume was used to generate 50 templates for template picking (270 Å diameter). The particles were subsequently extracted from the original micrographs with a box size of 784 px and Fourier-cropped to 392 px.

For each dataset, 2D classification, *ab initio* reconstruction (three classes, $C_1$ symmetry), and homogeneous refinement ($C_1$ symmetry) were performed, after which all the extracted particles were subjected to four rounds of heterogeneous refinement ($C_1$ symmetry), using 5 decoy classes and the homogeneously refined map as initial volumes. The particles from the best class resulting from the heterogeneous refinement were re-extracted using the full box size and homogeneously refined ($C_1$ symmetry) again. Afterwards, the particles were further sorted into two classes using 3D classification (filtered to 4 Å) before a final round of homogeneous refinement ($C_1$ symmetry) with local and global CTF refinement, per particle defocus refinement, and Ewald sphere correction. From all data sets, 65,020 particles were picked randomly for the last round of homogeneous refinement to ensure the comparability of the data. The global and local resolution was estimated using a gold-standard FSC threshold of 0.143. The structures were visualized with UCSF ChimeraX 1.9.[6,7]

**50S Ribosomal Subunit – Conventional**

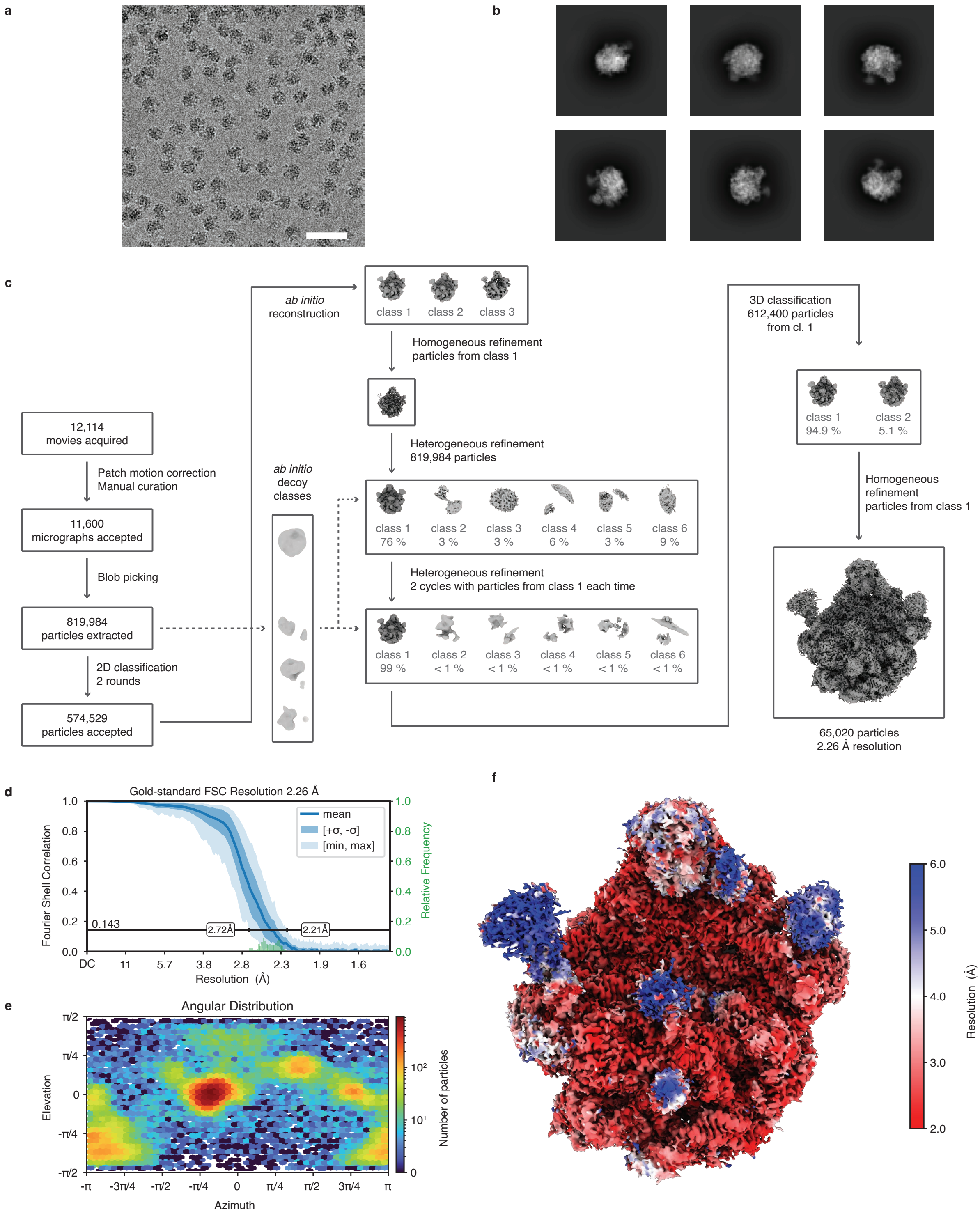

**Figure S3 | Cryo-EM processing workflow for the conventional cryo sample of the 50S ribosomal subunit. a,** Representative micrograph. Scale bar, 50 nm. **b,** 2D class averages gathered from the processing workflow. **c,** Data processing workflow in CryoSPARC. The symmetry applied is $C_1$ throughout the processing. The map is shown at a threshold of 3 σ above the mean. **d,** Conical FSC, with the cutoff of 0.143 indicated by a black line. The mean FSC is shown as a blue line, with the dark blue shading indicating one standard deviation, and the light blue shading the minimum and maximum values of the conical FSC. A histogram of the resolution values is shown in green, with the minimum and maximum values indicated. **e,** Orientation distribution of the particles. **f,** Final map with the local resolution estimation indicated in color (GSFSC cutoff of 0.5).

**50S Ribosomal Subunit – sealed and revitrified, 30 μs**

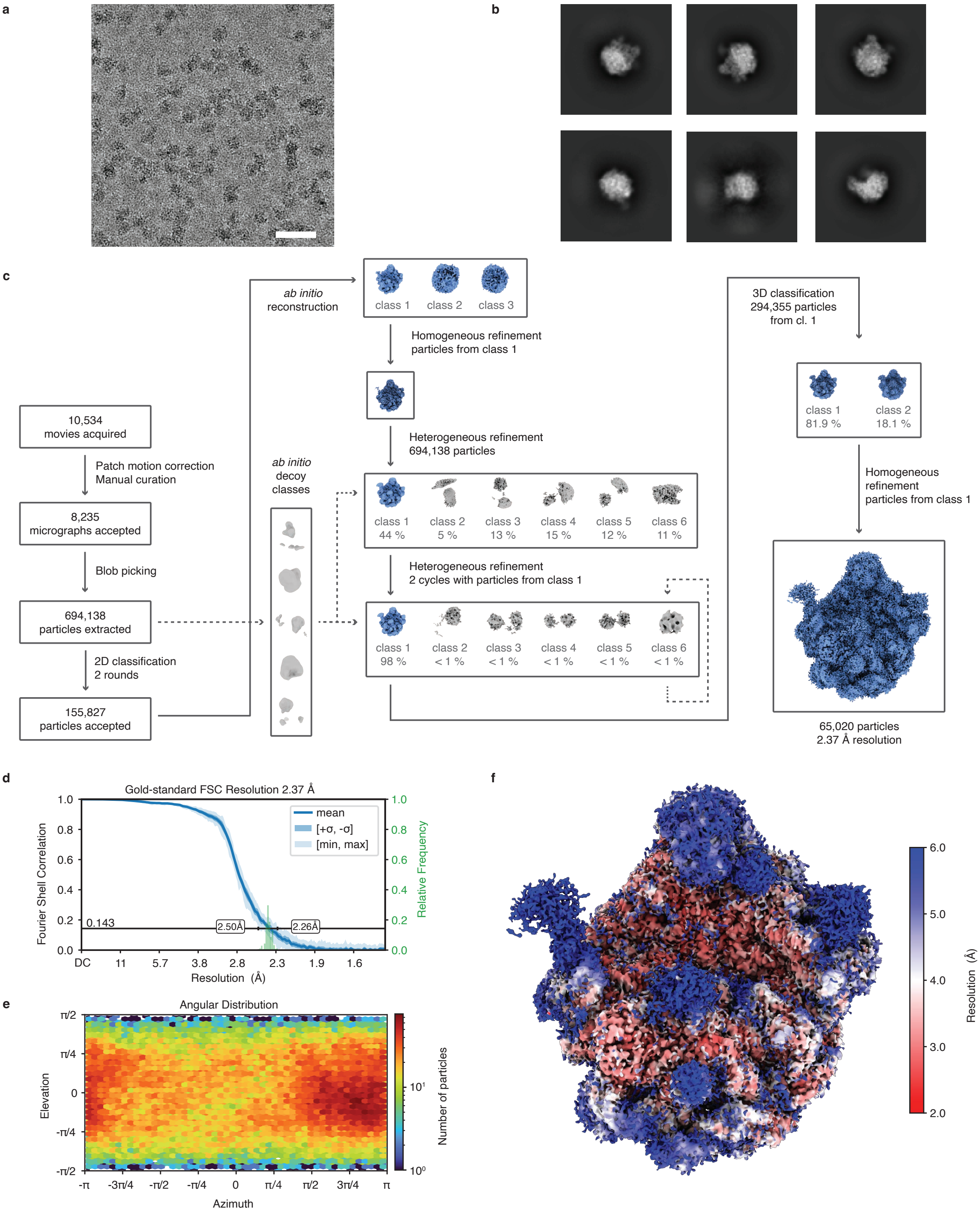

**Figure S4 | Cryo-EM processing workflow for the sealed and revitrified sample of the 50S ribosomal subunit (30 µs). a,** Representative micrograph. Scale bar, 50 nm. **b,** 2D class averages gathered from the processing workflow. **c,** Data processing workflow in CryoSPARC. The symmetry applied is $C_1$ throughout the processing. The map is shown at a threshold of 3 σ above the mean. **d,** Conical FSC, with the cutoff of 0.143 indicated by a black line. The mean FSC is shown as a blue line, with the dark blue shading indicating one standard deviation, and the light blue shading the minimum and maximum values of the conical FSC. A histogram of the resolution values is shown in green, with the minimum and maximum values indicated. **e,** Orientation distribution of the particles. **f,** Final map with the local resolution estimation indicated in color (GSFSC cutoff of 0.5).

**50S Ribosomal Subunit – sealed and revitrified, 150 µs**

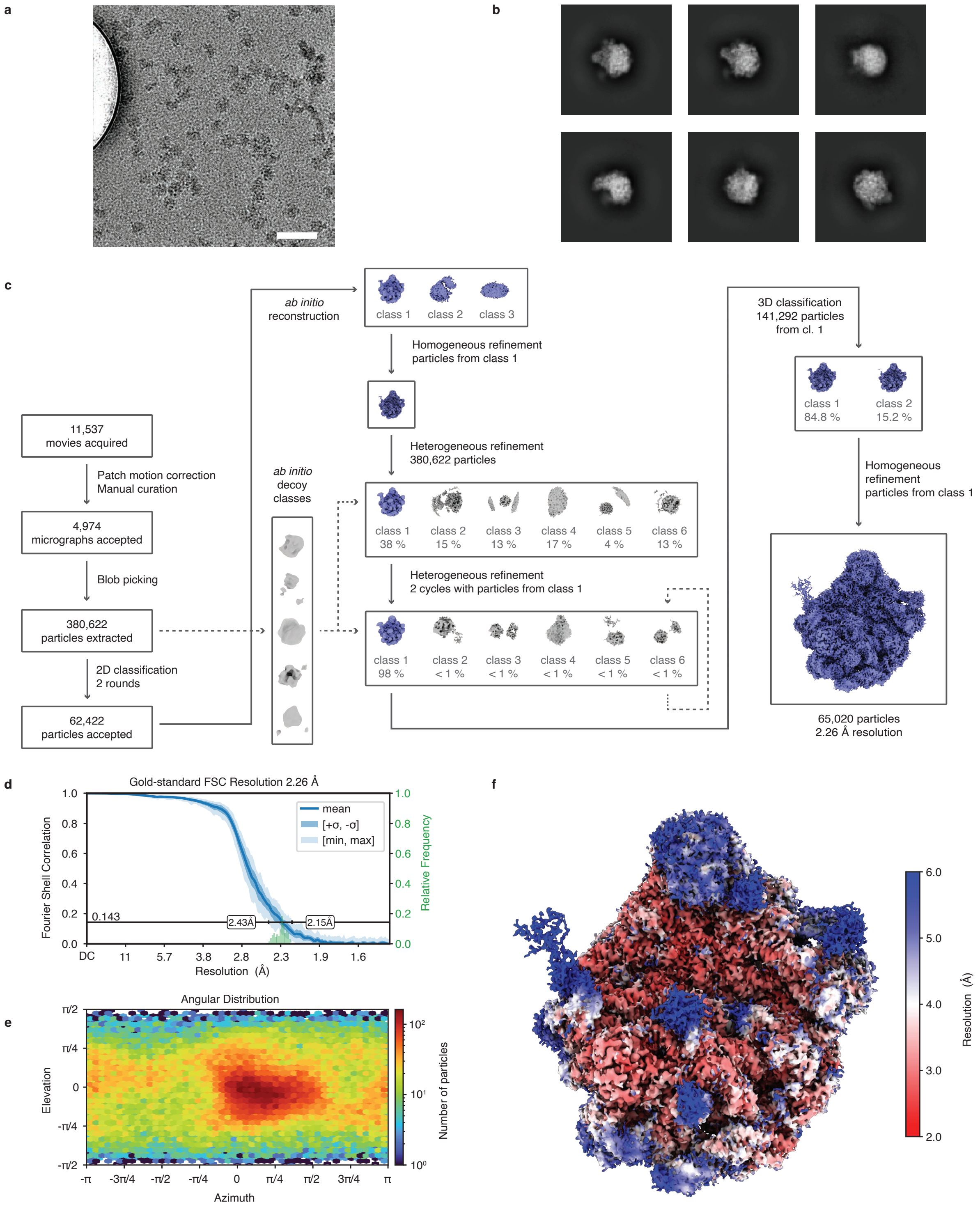

**Figure S5 | Cryo-EM processing workflow for the sealed and revitrified sample of the 50S ribosomal subunit (150 µs). a,** Representative micrograph. Bubble of gas formed between the silicon dioxide layers is visible on the left side of the image. Scale bar, 50 nm. **b,** 2D class averages gathered from the processing workflow. **c,** Data processing workflow in CryoSPARC. The symmetry applied is $C_1$ throughout the processing. The map is shown at a threshold of 3 σ above the mean. **d,** Conical FSC, with the cutoff of 0.143 indicated by a black line. The mean FSC is shown as a blue line, with the dark blue shading indicating one standard deviation, and the light blue shading the minimum and maximum values of the conical FSC. A histogram of the resolution values is shown in green, with the minimum and maximum values indicated. **e,** Orientation distribution of the particles. **f,** Final map with the local resolution estimation indicated in color (GSFSC cutoff of 0.5).

50S Ribosomal Subunit – sealed and revitrified, 300 μs

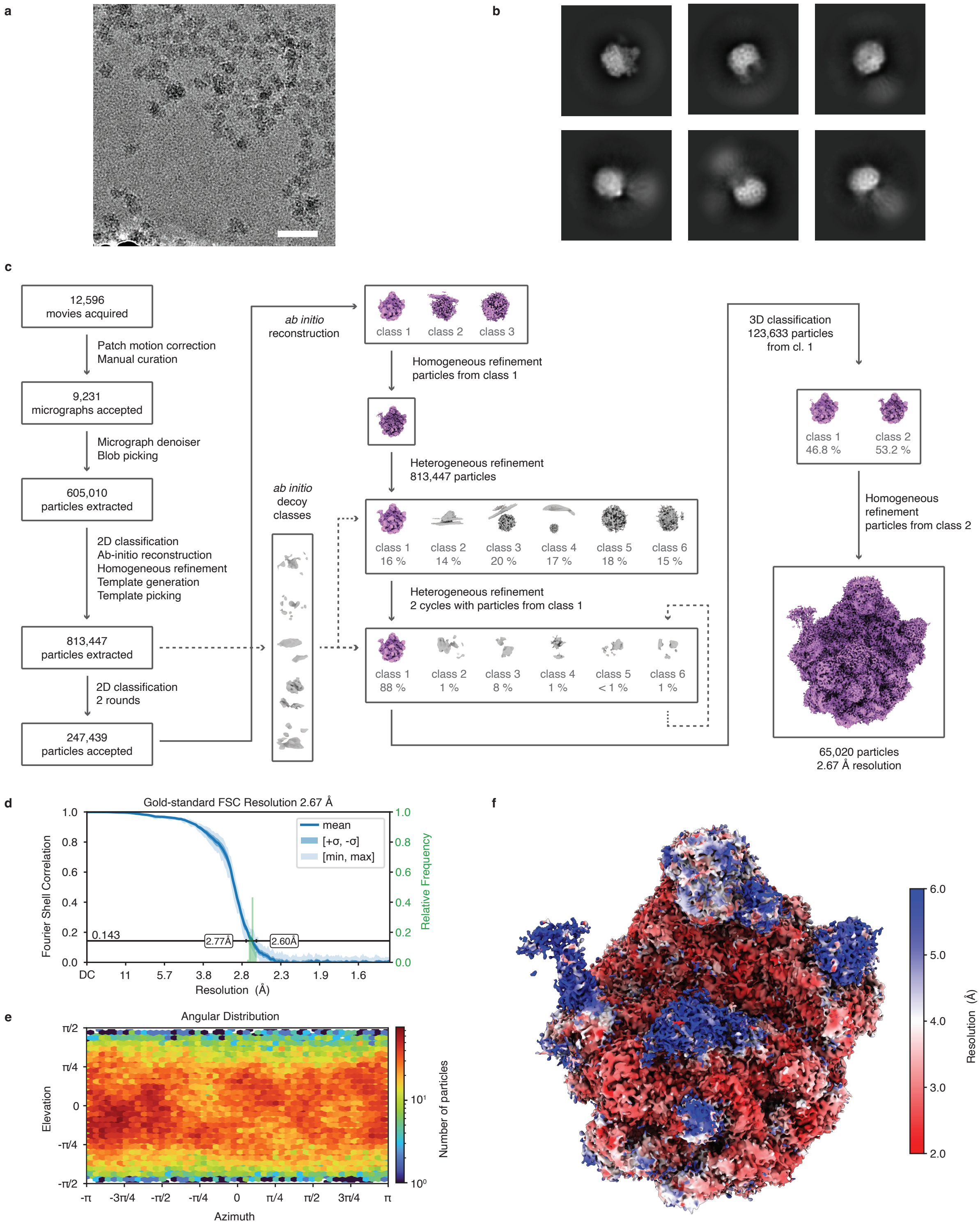

**Figure S6 | Cryo-EM processing workflow for the sealed and revitrified sample of the 50S ribosomal subunit (300 μs). a,** Representative micrograph. Scale bar, 50 nm. **b,** 2D class averages gathered from the processing workflow. **c,** Data processing workflow in CryoSPARC. The symmetry applied is $C_1$ throughout the processing. The map is shown at a threshold of 3 σ above the mean. **d,** Conical FSC, with the cutoff of 0.143 indicated by a black line. The mean FSC is shown as a blue line, with the dark blue shading indicating one standard deviation, and the light blue shading the minimum and maximum values of the conical FSC. A histogram of the resolution values is shown in green, with the minimum and maximum values indicated. **e,** Orientation distribution of the particles. **f,** Final map with the local resolution estimation indicated in color (GSFSC cutoff of 0.5).

**6 | Cryo-EM data collection and processing statistics**

| | **Apoferritin – conventional** | **Apoferritin – 210 μs** |
|---|---|---|
| ***Database Depositions*** | | |
| ***Collection and processing*** | | |
| *Microscope* | Thermo Fisher Titan Krios G4 | Thermo Fisher Titan Krios G4 |
| *Camera* | Falcon 4i | Falcon 4i |
| *Energy Filter (energy window)* | SelectrisX (10 eV) | SelectrisX (10 eV) |
| *Magnification* | 165,000 | 165,000 |
| *Accelerating Voltage (kV)* | 300 | 300 |
| *Exposure time (s/micrograph)* | 3.6 | 3.6 |
| *Exposure (e-/Å$^2$)* | 50 | 50 |
| *Defocus (μm)* | -0.5 to -1.2 | -0.5 to -1.2 |
| *Pixel size (Å/px)* | 0.732 | 0.732 |
| *Image format* | EER | EER |
| *Symmetry imposed* | O | O |
| *Initial number of particles* | 238,566 | 1,550,394 |
| *Final number of particles* | 170,473 | 170,473 |
| *Map resolution (Å)* | 1.66 | 1.79 |
| *FSC threshold* | 0.143 | 0.143 |
| *Map sharpening B factor (Å$^2$)* | 44.9 | 46.1 |
| *Orientation diagnostics* | | |
| *cFAR* | 0.93 | 0.93 |
| *SCF** | 0.970 | 0.979 |

**Table S1 |** Cryo-EM data collection and processing statistics for the mouse heavy-chain apoferritin.

| | 50S – conventional | 50S – 30 µs | 50S – 150 µs | 50S – 300 µs |
|---|---|---|---|---|
| ***Database Depositions*** | | | | |
| ***Collection and processing*** | | | | |
| *Microscope* | Thermo Fisher Titan Krios G4 | Thermo Fisher Titan Krios G4 | Thermo Fisher Titan Krios G4 | Thermo Fisher Titan Krios G4 |
| *Camera* | Falcon 4i | Falcon 4i | Falcon 4i | Falcon 4i |
| *Energy Filter (energy window)* | SelectrisX (10 eV) | SelectrisX (10 eV) | SelectrisX (10 eV) | SelectrisX (10 eV) |
| *Magnification* | 165,000 | 165,000 | 165,000 | 165,000 |
| *Accelerating Voltage (kV)* | 300 | 300 | 300 | 300 |
| *Exposure time (s/micrograph)* | 3.6 | 3.6 | 3.6 | 3.6 |
| *Exposure (e-/Å$^2$)* | 50 | 50 | 50 | 50 |
| *Defocus (µm)* | -0.5 to -1.2 | -0.5 to -1.2 | -0.5 to -1.2 | -0.5 to -1.2 |
| *Pixel size (Å/px)* | 0.732 | 0.732 | 0.732 | 0.732 |
| *Image format* | EER | EER | EER | EER |
| *Symmetry imposed* | $C_1$ | $C_1$ | $C_1$ | $C_1$ |
| *Initial number of particles* | 819,984 | 694,138 | 380,622 | 813,446 |
| *Final number of particles* | 65,020 | 65,020 | 65,020 | 65,020 |
| *Map resolution (Å)* | 2.47 | 2.37 | 2.26 | 2.67 |
| *FSC threshold* | 0.143 | 0.143 | 0.143 | 0.143 |
| *Map sharpening B factor (Å$^2$)* | 34.0 | 30.9 | 29.7 | 37.2 |
| *Orientation diagnostics* | | | | |
| *cFAR* | 0.49 | 0.88 | 0.76 | 0.88 |
| *SCF** | 0.526 | 0.987 | 0.942 | 0.989 |

**Table S2 |** Cryo-EM data collection and processing statistics for the 50S ribosomal subunit.

**7 | Irradiation with long laser pulses and breakup of the thin liquid film**

To reach long timescales, we melt the sample several times with laser pulses of 30 µs duration instead of using a single long laser pulse. This is advantageous because under laser irradiation, the thin liquid film breaks up more frequently as the pulse duration is increased. This is illustrated in Fig. S7, which shows micrographs of typical samples after irradiation with long laser pulses. Flash melting with a single 100 µs laser pulse often leaves the sample intact, as shown in Fig. S7a. In the center of the laser focus, the sample has revitrified (red circle), while adjacent areas have crystallized. A diffraction pattern collected from one of the central holes confirms the presence of vitreous ice (inset). In contrast, if we irradiate a similar area with a 200 µs laser pulse of the same power, the thin liquid film usually breaks up, as shown in Fig. S7b. While crystallization has occurred outside of the area marked with a red circle, holes within this area are mostly empty, even though they are still covered with silicon dioxide membranes. A diffraction pattern collected from one of the central holes shows that no vitreous ice is left (inset) and only a diffuse signal from amorphous silicon dioxide remains. For the experiments in Fig. S7, the laser power was adjusted such that a 30 µs laser pulse can successfully revitrify a similar sample area. During a 100 µs laser pulse, the sample reaches the same temperature. This can be seen in the heat transfer simulations of Fig. S12b, which show that under laser irradiation, the sample temperature initially rises rapidly before it plateaus after a few microseconds. After that, the temperature remains practically constant until the laser is switched off.

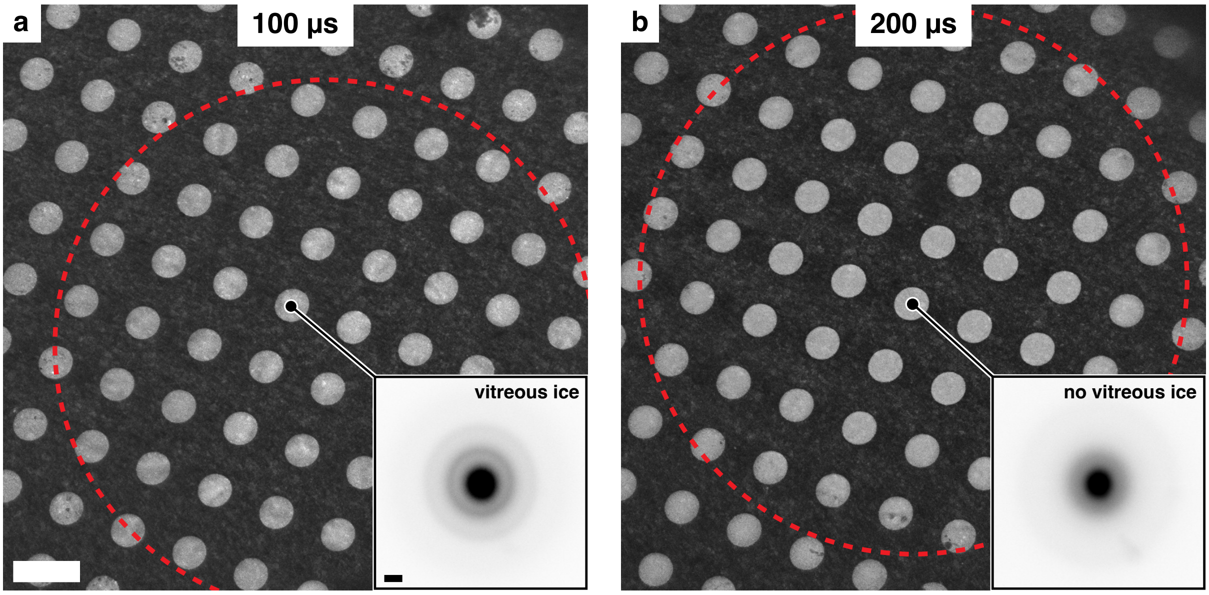


**Figure S7 | With increasing laser pulse duration, breakup of the thin liquid film becomes more frequent. a,** Micrograph of a sample after flash melting with a 100 µs laser pulse, with the revitrified region marked by a red circle. A diffraction pattern confirms the presence of vitreous ice (inset). **b,** Irradiation of a similar area with a 200 µs laser pulse causes the sample to break up. Most holes within the area marked by the red circle are empty even though they are covered by silicon dioxide membranes, as confirmed by the diffraction pattern in the inset. Scale bars, 2.5 µm and 1 Å$^{-1}$.

**8 | Variability analysis of the L1 stalk motion of the 50S ribosomal subunit**

In the temperature jump experiment of Fig. 5, we analyze the conformational distribution of the L1 stalk of the 50S ribosomal subunit with the help of a 3D variability analysis in CryoSPARC.[5] The corresponding workflow is illustrated in Fig. S8. First, the volumes obtained for the three transient ensembles (flash melting for 30 µs, 150 µs, and 300 µs, Fig. S4-6) are aligned to a common reference, for which we use the reconstruction from the conventional cryo sample (Fig. S3). After Fourier cropping to a box size of 392 pixels, the particles of the transient ensembles are used for a joint reconstruction, which is then subjected to non-uniform refinement with dynamic masking disabled. Subsequently, each of the three transient ensembles is separately orientation rebalanced (192 orientation bins, rebalanced percentile 80, random intra-bin exclusion criterion) to prevent the variability analysis from being biased towards motions that appear prominently in the projections associated with preferred particle orientations. Finally, we perform a 3D variability analysis of the joint dataset (resolution filter of 10 Å, 20 iterations) with a static mask that isolates the L1 stalk motion. The mask was built with ChimeraX[6,7] from the initial reconstruction of the joint dataset (Gaussian filtered with $\sigma = 2.5$ Å) by isolating the region surrounding the L1 stalk and applying a 10 pixel dilation (12 pixel soft padding) to the volume in CryoSPARC. The resulting mask is shown in Figure S8. Figure S9, illustrates the first two components of the variability analysis, which correspond to approximately orthogonal wagging motions of the L1 stalk.

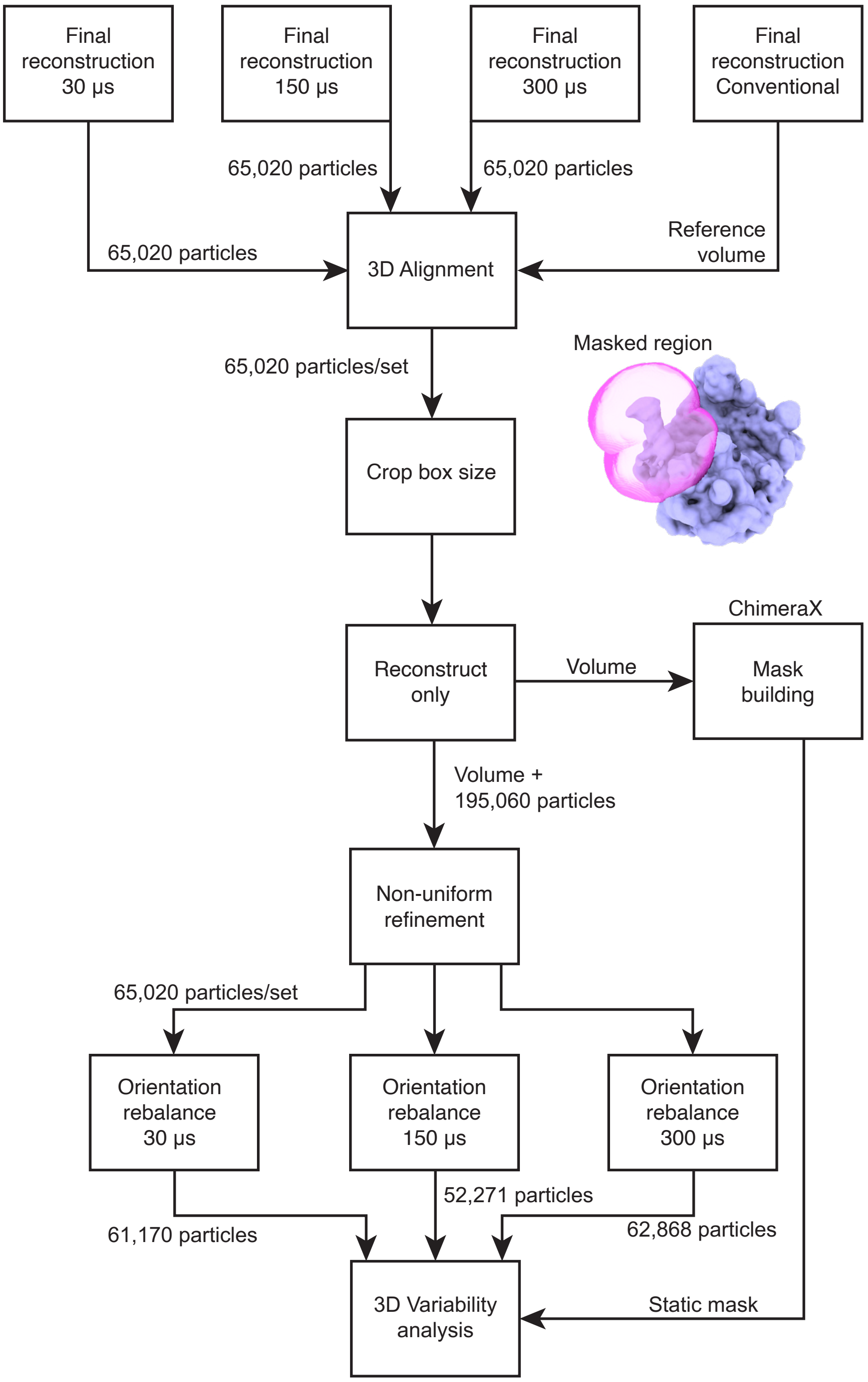


**Figure S8 | Workflow for the 3D variability analysis of the L1 stalk motion of the 50S Ribosomal subunit.**

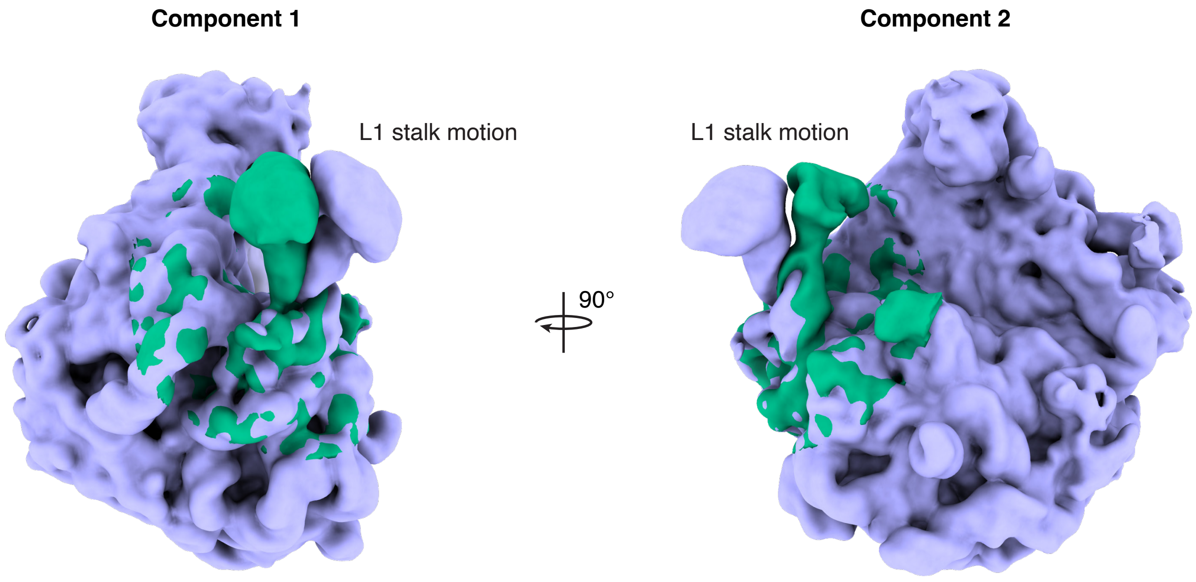


**Figure S9 | Illustration of the first two principal components of the L1 stalk motion of the 50S ribosomal subunit as obtained from a 3D variability analysis.** The purple and green volumes represent reconstructions from opposite wings of the conformational distribution (Gaussian filtered to 2.5 Å), with the green volume overlayed in the region that was included in the variability analysis (Fig. S8).

**9 | Determination of the conformational distributions for the L1 stalk movement**

In the experiment presented in Fig. 5, we show that the amplitude of the L1 stalk motion of the 50S ribosomal subunit responds to a temperature jump on a timescale of hundreds of microseconds. As a measure of the amplitude of motion, we use the standard deviation of the conformational distribution of the L1 stalk along its first principal component of motion (Fig. 5a, Supplementary Information 8). Obtaining an accurate conformational distribution for this wagging motion faces the challenge that in some particle orientations, the projected amplitude of this motion is so small that it becomes difficult for the algorithm to accurately assign a value for this principal component. The consequence of this geometric effect can be seen in Fig. S10a, which shows the angular distribution of the apparent width of the conformational distribution (as obtained from a variability analysis). The angular distribution shows two maxima (marked 1 and 2), incorrectly suggesting that particles with these orientations exhibit a larger amplitude of motion. Figure S10b reveals that when viewed along these directions, the wagging motion of the L1 stalk is obscured. Apparently, this results in a large uncertainty of the assignment of the particles to a value of the principal component and thus artificially broadens the extracted conformational distribution.

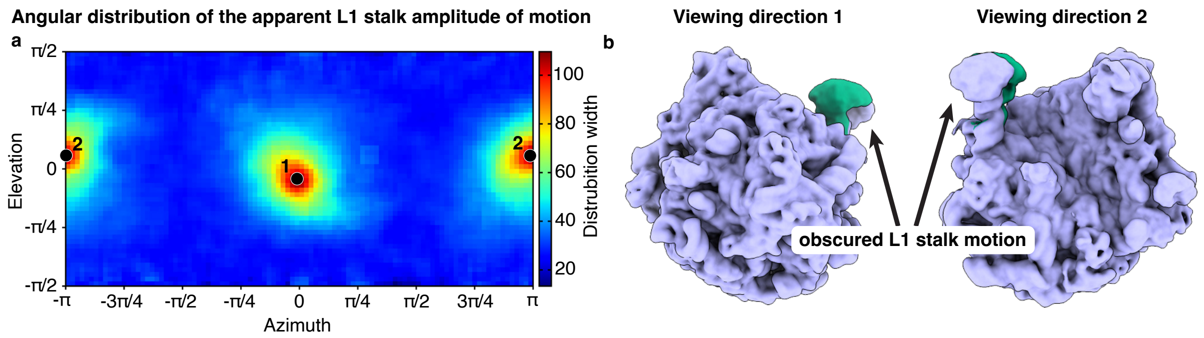


**Figure S10 | Particle orientation affects the accuracy with which conformational distributions can be determined. a,** Angular distribution of the apparent width of the conformational distribution (as measured by the standard deviation) for the first component of the L1 stalk movement. Two particle orientations (marked 1 and 2) appear to show a significantly wider distribution. The distribution is shown for a mixed dataset of the three transient ensembles. Each pixel represents the width of the conformational distribution for the particles contained in the surrounding 5 × 5 pixels. **b,** When viewed along the directions corresponding to the maxima in **a**, the wagging motion of the L1 stalk is obscured. This apparently results in a larger uncertainty for the assignment of the particles to a value of the principal component, which artificially broadens the extracted conformational distribution.

The subtle changes of the width of the conformational distribution in response to the temperature jump are obscured by the artificial broadening of the distribution due to particle orientations for which the L1 stalk motion is hidden. We therefore remove particle orientations in which the projected amplitude of motion is small, as illustrated in Fig. S11. Figure S11a shows the angular distributions of the apparent width of the conformational distributions (as measured by the standard deviation) for the first two variability components. These distributions bear close resemblance to the relative projected amplitudes of motion shown in Fig. S11b. The projected amplitudes of motion are derived from the difference of two volumes in which the L1 stalk is displaced by equal amounts in opposite directions along a variability component. We then apply the mask to this difference volume that we have used in the variability analysis and project it along all viewing directions. Finally, we calculate the sum of the absolutes of all pixels for each of these two-dimensional projections to obtain the projected amplitudes of motion, which we then normalize. We remove particle orientations from our analysis for which the relative projected amplitude of motion is below a threshold of 0.6. This is illustrated in Fig. S11c, which shows the angular distributions of the apparent width of the conformational distributions with the excluded orientations in grey. We note that varying this threshold within a wide range does not qualitatively change our results. This is particularly the case for the 30 µs and 300 µs datasets, which are of somewhat higher quality than the 150 µs dataset.

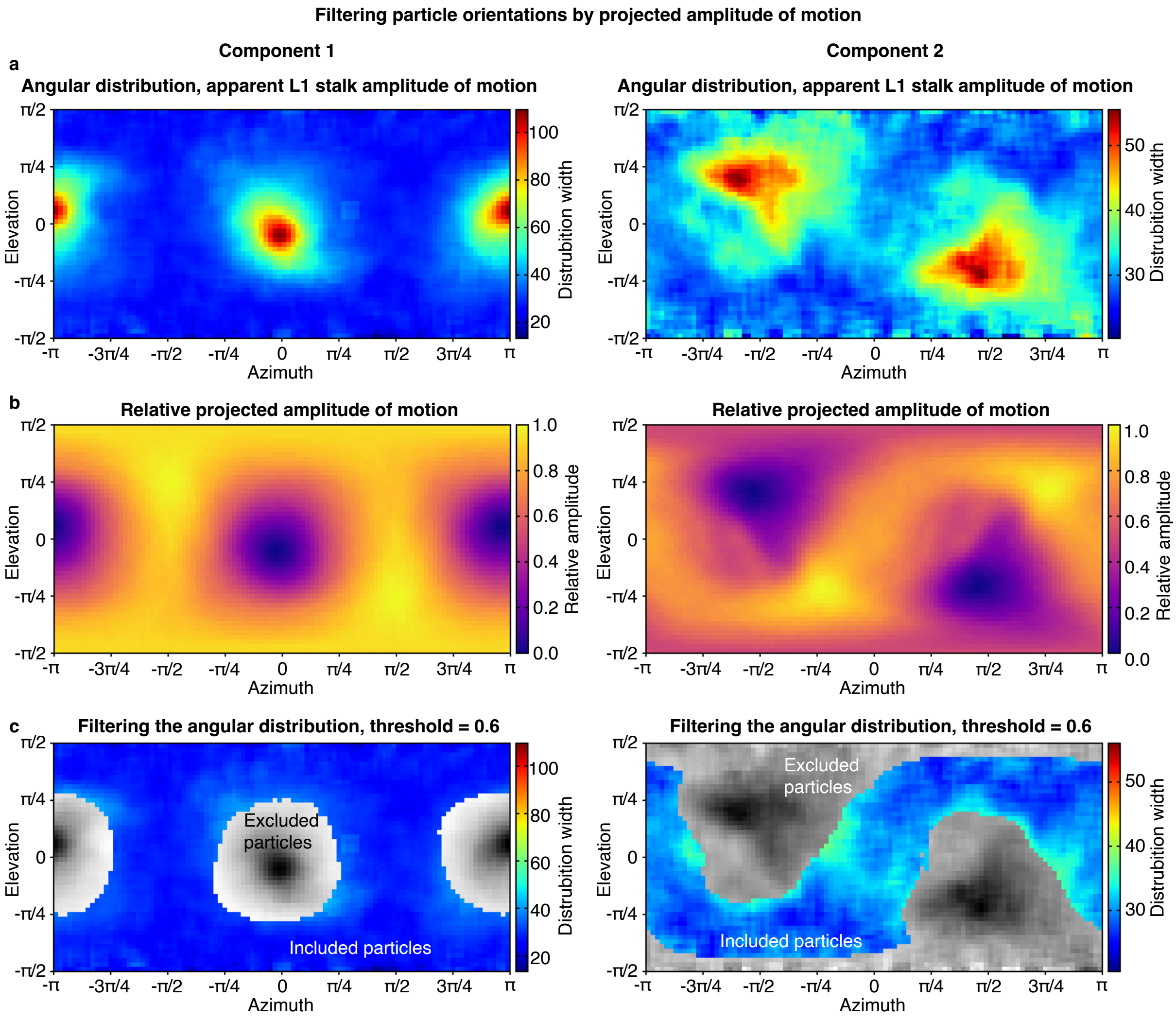


**Figure S11 | Filtering the particle orientations by projected amplitude of motion. a,** Angular distribution of the apparent width of the conformational distribution, as measured by the standard deviation, for the first two variability components. Each pixel represents the width of the conformational distribution for the particles contained in the surrounding 5 × 5 pixels. **b,** Angular distribution of the relative projected amplitude of motion. **c,** To obtain accurate conformational distributions, we remove particle orientations (grey) for which the relative projected amplitude of motion in **b** is below a threshold of 0.6.

### 10 | Simulation of the sample temperature during laser heating

We simulate the temperature evolution of the sample during laser heating in COMSOL Multiphysics 6.3 as previously described,[3,8] with only small modifications of the simulation geometry to match the experiments described here. We simulate a typical ice thickness of 50 nm within the grid holes and 25 nm on top of the gold foil. To reduce computational cost, we omit the thin silicon dioxide sealing membranes, whose heat capacity and thermal conductivity are so small that they can be neglected. Moreover, we assume that the sample cannot evaporate and therefore remove evaporative cooling from our model. The sample is heated with a Gaussian laser beam (22 µm FWHM spot size in the sample plane) that is aimed at the center of a grid square. We vary the simulated laser power such that the resulting diameter of the revitrified area covers the range of diameters that we observed in our experiment. Figure S12a,b shows a typical temperature evolution of the sample in the center of the laser focus under illumination with a 30 and 100 µs laser pulse, respectively. After the laser is switched on, the sample temperature rises rapidly and crosses the melting point at about 7.5 µs. After 15 µs, the temperature plateaus at approximately 302 K and remains stable for the duration of the laser pulse. After the end of the laser pulse, the sample cools rapidly, dropping to the glass transition temperature of 136 K after about 4 µs. Note that while the heating and cooling times increase slightly with increasing ice thickness, the temperature at which the sample stabilizes barely changes within the small range of ice thicknesses in our experiment.[4]

Figure S12c shows a typical temperature distribution of the sample at the end of a 30 µs laser pulse, with the 273 K isotherm indicated, which approximately marks the size of the area that is revitrified by the laser pulse (from Fig. 5b). Note that the sample reaches this temperature distribution within about 15 µs during laser irradiation, after which the temperature experienced by a particle in a given location remains approximately constant until the laser is switched off. Figure S12d shows simulated temperature profiles (thin lines) for different laser powers together with fifth-order polynomial fits (bold). With increasing laser power, the sample reaches higher temperature, and the diameter of the revitrified area grows.

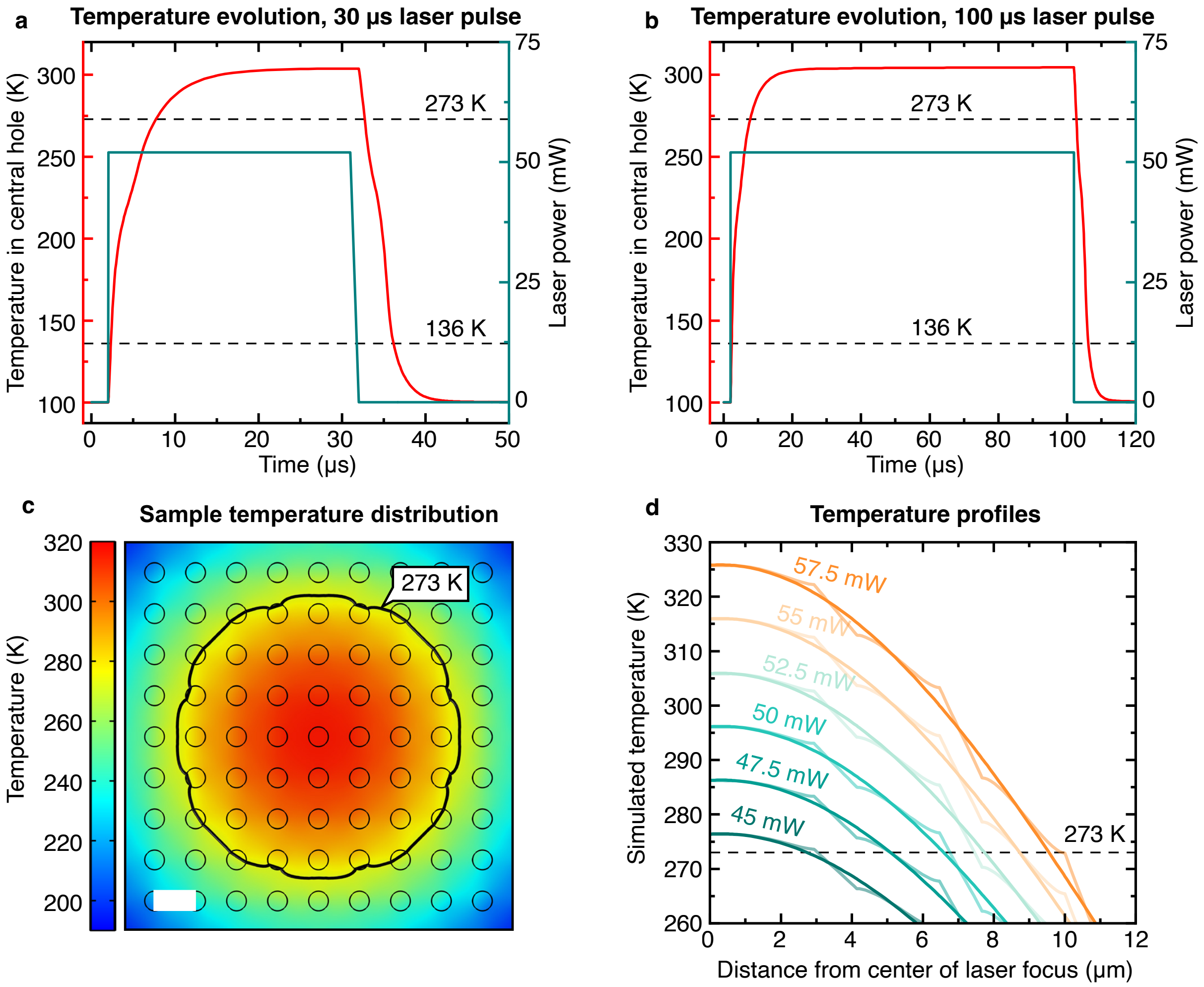


**Figure S12 | Simulation of the temperature evolution of the sample. a,b,** Temperature evolution of the sample within the hole in the center of the laser focus under illumination with a 30 and 100 µs laser pulse, respectively. **c,** Simulated temperature distribution of the sample at the end of a 30 µs laser pulse under typical conditions (from Fig. 5b). The black line indicates the 273 K isotherm, which approximately encloses the revitrified region. Scale bar, 2.5 µm. **d,** Simulated temperature profiles for different laser powers (thin lines) together with fifth-order polynomial fits (bold). The temperature profiles are taken diagonally across the grid square. The indicated temperature is that of the cryo sample, just above the holey gold support.

**11 | Estimate of the particle temperature during laser heating**

We estimate the temperature of the particles during laser heating with the help of the simulations presented in Supplementary Information 10. They show that the temperature of each particle can be approximately determined if the radius of the revitrified area is known as well as distance of the particle from its center. We obtain both quantities as follows. During high-resolution data collection, we attempt to acquire micrographs across the entire revitrified area, with some micrographs also collected in the surrounding regions that have crystallized. The micrographs of the revitrified area form a circular patch that is surrounded by micrographs exhibiting crystallization. We can then approximately determine the center and radius of the revitrified area from the smallest circle that encloses the center coordinates of all micrographs with vitreous ice (as calculated from the corresponding stage positions and beam shifts). This determination is more difficult if we have not been able to completely map out the revitrified area with high-magnification micrographs, for example, because it is partially covered with ice contaminations. We therefore additionally consider a low-magnification micrograph of each grid square, in which the center of the revitrified area appears as a circle of lighter contrast. With both pieces of information together, we can then determine the center and radius of the circle that encloses the revitrified area, a task that can be accomplished with sufficient accuracy by eye. Revitrified areas for which this method does not provide a reliable estimate are rejected. To obtain the radius of the revitrified area, we increase the radius of the smallest circle enclosing all the micrographs centers by half the width of a micrograph (150 nm). We approximate the distance of each particle from the center of the revitrified area by the distance of the center of the corresponding micrograph from the center of the enclosing circle.

Finally, we estimate the temperature of each particle as follows. The fifth-order polynomial fits of the simulated temperature profiles in Fig. S12d approximate a surface that describes the particle temperature as a function of the radius of the revitrified area and the distance of the particle from its center. With estimates of both quantities in hand, we then perform a linear interpolation of this surface to obtain the approximate temperature for each particle.

### 12 | Evolution of the amplitude of the L1 stalk motion

We use the following procedure to characterize how the amplitude of the L1 stalk motion evolves in the temperature jump experiment of Fig. 5. As detailed in Supplementary Information 8, we determine the principal components of the L1 stalk motion from a variability analysis of the merged datasets of the transient ensembles (30 μs, 150 μs, and 300 μs of laser melting). The first two components correspond to wagging modes (Fig. S9). For both, we filter the datasets to remove particle orientations that have a large uncertainty for assigning the value of the principal component (Supplementary Information 9). The temperature of each particle during laser melting is then estimated from simulations (Supplementary Information 11). Finally, we bin the particles by their estimated temperature and determine the widths of the conformational distributions in each bin, as measured by the standard deviation. Figure S13 displays the distribution width as a function of the simulated particle temperature. We obtain a qualitatively similar result for both wagging motions of the L1 stalk. After 30 μs, the distribution width is largely independent of temperature (blue), indicating that the conformational ensemble still mostly reflects the temperature at which the sample was initially prepared (293 K). The conformational distribution begins to narrow in the colder parts of the sample at 150 μs (orange). Finally, a pronounced temperature dependence is observed after 300 μs (purple). Note that at 30 μs, colder particles appear to have slightly wider conformational distributions. This trend may arise because the sample is thicker in the colder parts of the revitirified area, which increases the uncertainty for assigning the principal component and thus artificially increases the width of the distribution.

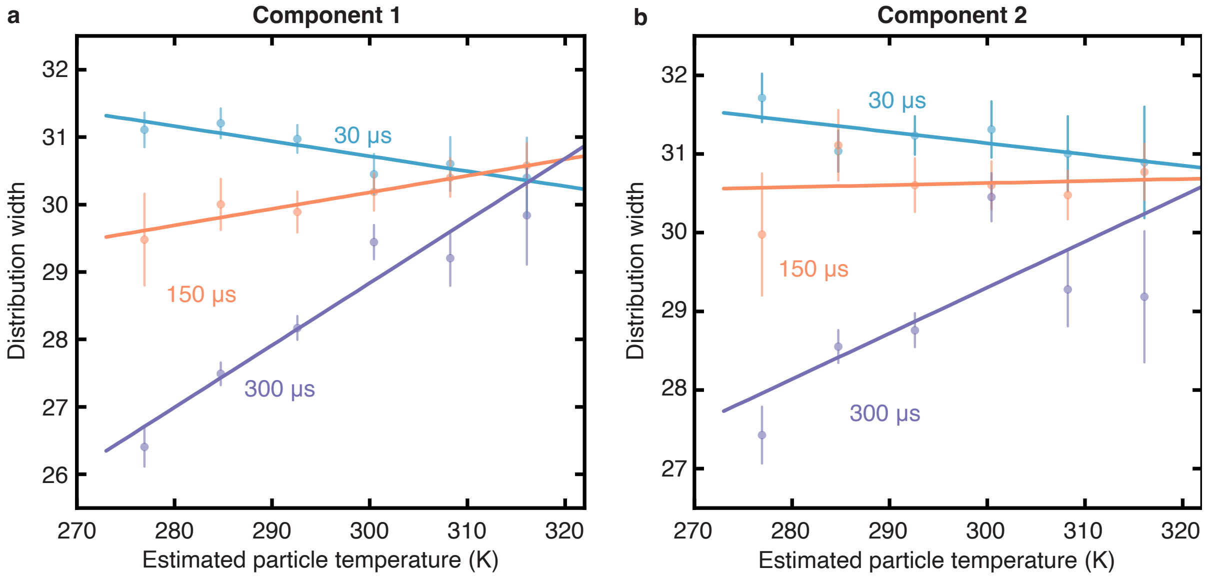


**Figure S13 | Dynamics of the 50S ribosomal subunit in a microsecond time-resolved temperature jump experiment.** The width of the conformational distribution along the first two principal components (Fig. S9, as measured by the standard deviation) is displayed as a function of the estimated particle temperature after 30 µs (blue), 150 µs (orange), and 300 µs (purple) of laser melting. Error bars represent standard error of the standard deviation.[9] Linear fits of the data are added as a guide to the eye.
